# Dynamics of spontaneous alpha activity correlate with language ability in young children

**DOI:** 10.1101/443192

**Authors:** Elaine Y. L. Kwok, Janis Oram Cardy, Brian L. Allman, Prudence Allen, Björn Herrmann

**Author notes:** Corresponding Authors: Elaine Kwok, Communication Sciences and Disorders, The University of Western Ontario, London, ON, Canada N6G 1H,; Björn Herrmann, Brain and Mind Institute, The University of Western Ontario, London, ON, Canada N6A 5B7. Declaration of interest: None.

## Abstract

Early childhood is a period of tremendous growth in both language ability and brain maturation. To understand the dynamic interplay between neural activity and spoken language development, we used resting-state EEG recordings to explore the relation between alpha oscillations (7–10 Hz) and oral language ability in 4- to 6-year-old children with typical development (*N*=41). Three properties of alpha oscillations were investigated: a) alpha power using spectral analysis, b) flexibility of the alpha frequency quantified via the oscillation’s moment-to-moment fluctuations, and c) scaling behavior of the alpha oscillator investigated via the long-range temporal correlation in the alpha-amplitude time course. All three properties of the alpha oscillator correlated with children’s oral language abilities. Higher language scores were correlated with lower alpha power, greater flexibility of the alpha frequency, and longer temporal correlations in the alpha-amplitude time course. Our findings demonstrate a cognitive role of several properties of the alpha oscillator that has largely been overlooked in the literature.

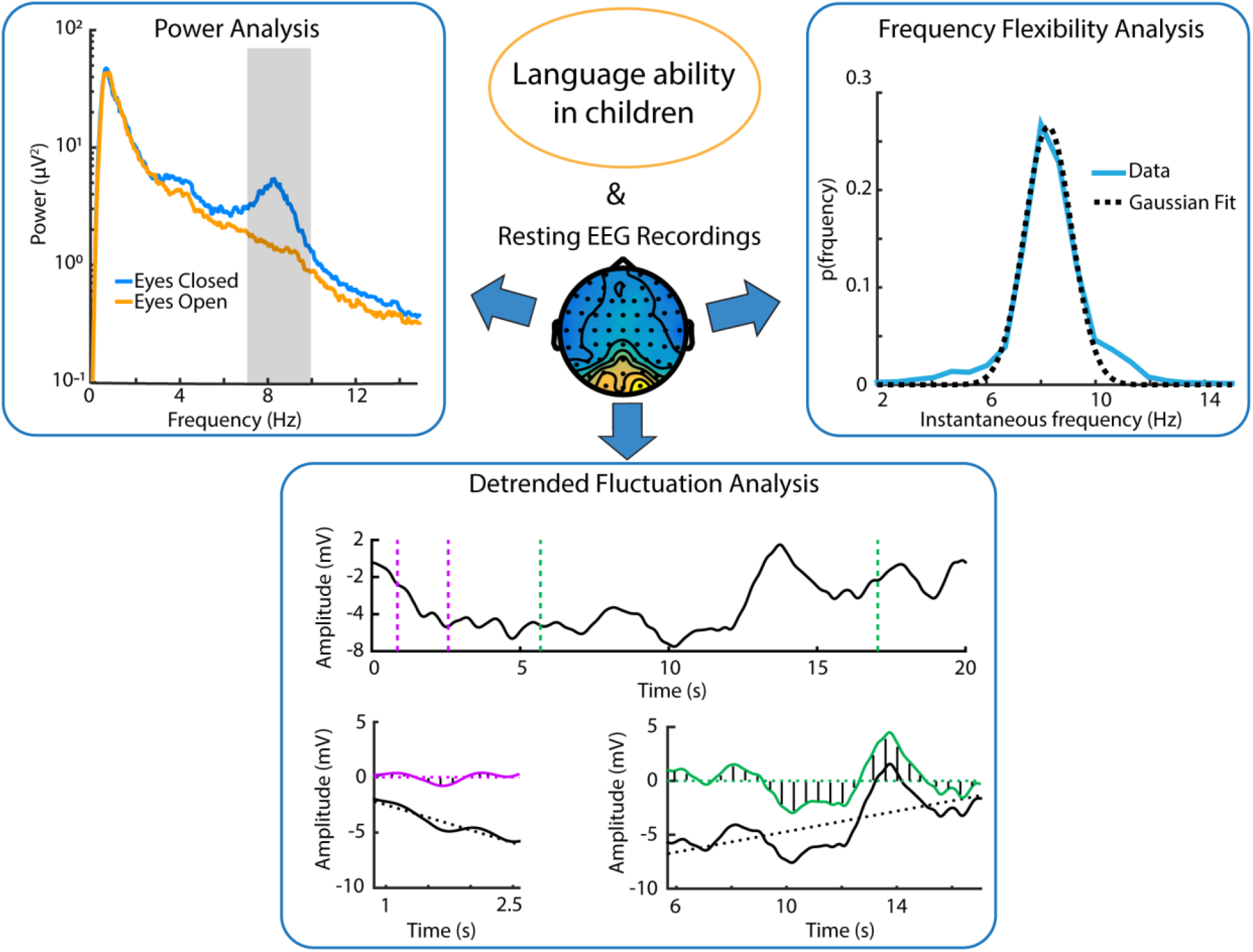
Graphical Abstract.

## 1. Introduction

In the first six years of life, children show tremendous growth in their spoken language development, moving from first words to combining a few words at a time with limited grammar, to speaking in increasingly complex and adult-like sentences [1–5]. At the same time, children’s neural and perceptual sensitivity to sounds improves and they become better at discriminating different frequencies, intensities, and durations of auditory stimuli [6,7]. This maturation of auditory perceptual abilities has important implications for oral language development [8]. This early period is also a time during which children may be first identified as having *developmental language disorder* (also known as specific language impairment; 9,10) and other neurodevelopmental disorders that include impaired language development (e.g., autism spectrum disorder, intellectual disability). Individual differences in neural function that underlie oral language development are still being investigated, and a reliable neural marker for language disorder has not been identified.

Spontaneous neural oscillations, as recorded using electroencephalography (EEG) when individuals are at rest, may be a particularly promising tool for understanding the neural underpinnings of language development and disorders. They can be recorded without any sensory stimulation (as opposed to measuring event-related potentials), making this a useful tool for young children or clinical populations who struggle to participate in or tolerate paradigms for collecting event-related potentials [11]. The dominant spontaneous oscillation found in the human EEG is alpha activity, which has repeatedly been linked to cognition, particularly in relation to cortical inhibition and attention [12–15]. Studies of individuals with disorders involving written language report a relation between spontaneous alpha power and reading ability [16–20], however, the role of spontaneous alpha activity in oral language ability has not been investigated. Given that the many cognitive roles of alpha oscillations and the connection between oral and written language skill development [21,22], we might also expect to find a relation between the power of spontaneous alpha activity and the ability to comprehend or use spoken language. It is important to recognize that while spontaneous frontal gamma activity (31–50 Hz) has also been found to associate with concurrent oral language skills [23] and later language skills [24], the low signal-to-noise ratio nature of spontaneous gamma activity poses challenges to some analytical approaches [25], making it a less than ideal candidate for the current study.

In addition to exploring the role of spontaneous alpha oscillation power in oral language ability, the current study aims to broaden the understanding of neural oscillations in language ability by employing multiple EEG analysis methodologies. The analysis of neural oscillations has relied almost exclusively on the investigation of the EEG power spectrum in the literature on both written [17,19,24, 25] and spoken language development/disorders [21,22], or on spatial coherence of activity in different frequency bands [28–30] (also see reviews [31,32]). While these traditional EEG analyses provide insights about neural energy (i.e., power) in specific frequency bands of the EEG, they fail to reveal other properties of EEG oscillatory activity.

Recent approaches to the study of neural oscillations emphasize that oscillatory activity may be non-sinusoidal in nature [33–36] and that non-sinusoidal properties of neural oscillators are related to neural communication and coordination [33,34]. Non-sinusoidal oscillations may appear as intermittent short bursts of energy in continuous EEG recordings [34,37,38]. Depending on the flexibility of the underlying neural oscillator, neural activity bursts will occur very regularly (i.e., at a fixed frequency) or occur at irregular points in time (i.e., the oscillator’s frequency varies over time). Estimation of the range of frequencies an oscillator expresses over time may thus provide information about the oscillator’s flexibility (as opposed to rigidity, that is, a fixed frequency).

A growing body of literature has also found that arrhythmic electrophysiological activities without a characteristic temporal periodicity (referred to as *scale-free dynamics*) predominate in the time series of human resting-state EEG [39–42]. Those dynamics are characterized by a power-law structure in the power spectrum (1/*f*, for example, amplitudes decrease as a function of frequency in an EEG power spectrum), which is present both across a broad frequency range [35,40] and within narrow frequency bands [43–46]. In the context of neural activity, 1/*f* structure may be indicative of self-organization, where amplitude fluctuations at any given moment can influence the amplitude of upcoming oscillations (known as *temporal memory effects*, [47]). The specific 1/*f* structure of a system is related to the temporal organization of its time series. One way to characterise the temporal organization of neural activity (and thus the 1/*f* scaling behavior) is through estimating the long-range correlation (or auto-correlation) in time courses of neural activity (e.g., using detrended fluctuation analysis; [25,48]). This temporal organization of the brain may play significant developmental and cognitive roles. Studies have revealed significant maturational changes in the temporal organization of EEG activities from childhood to adolescence (also continuing into adulthood; [49]) and that the temporal correlations are modulated by task-demand (e.g. eyes opening and closing: [50]; cued-button press: [39]). Nevertheless, the precise cognitive role of this temporal organization is currently unclear.

In this study, we explored the relation between oral language ability during early childhood and several measures of resting-state alpha activity: spectral power, flexibility of alpha frequency, and long-range temporal correlation in the alpha-amplitude time series.

## 2. Methods and Materials

### 2.1 Participants

Forty-one children with typical development aged 4–6 years participated in the current study (4 years, *n* = 12; 5 years, *n* = 14; 6 years, *n* = 15; median age = 4.58, 5.21, 6.58 years, respectively). Participants were recruited either as part of an epidemiological investigation of the language, reading and arithmetic skills of 4- to 10-year-old children in a local school board [51] or through personal contacts. Participants had no neurological, hearing, or visual impairments by parental report. To be included in the study, children had to score within the normal range (i.e., higher than −1*SD* from mean) on the Performance IQ (PIQ) of the *Wechsler Abbreviated Intelligence Scale* (WASI; [52]) and Core Language Score (CLS) of the *Clinical Evaluation of Language Fundamentals–Preschool - 2* (CELF-P2; [53]). We used standard scores from the PIQ and CLS for our analyses because they estimate the relative strength of each participant’s abilities compared to a population of same-age peers, that is, they are norm-referenced and age-corrected. Participants’ standard scores on the CLS and PIQ ranged from 86-119 (*M* = 106, *SD* = 9) and 88-131 (*M* = 109, *SD* = 12), respectively. All participants completed the language and performance IQ testing on the same day as their EEG recording.

### 2.2. EEG recording

Resting state EEG was recorded at a 500-Hz sampling frequency using a 128-channel DenseArray EEG system with HydroCel Geodesic Sensor Nets (Electrical Geodesics Inc., Eugene, OR, USA). An online hardware Bessel filter with a high-pass of 0.1 Hz and a low-pass of 100 Hz was applied to the recorded data. Participants were instructed to remain still and keep their eyes open or closed for one minute at a time. Three repetitions of alternating eyes-open and eyes-closed trials were conducted, resulting in a total of three minutes of EEG recording for both the eyes-open and the eyes-closed conditions.

### 2.3 EEG preprocessing

Data were analyzed offline using the fieldtrip toolbox in Matlab software [54] and custom Matlab scripts. Line noise was suppressed using a 60 Hz elliptic notch filter (cut-off frequencies: 59.2 Hz and 60.8 Hz; 70 dB suppression). EEG recordings were high-pass filtered (0.4 Hz, 2091 points, Hann window, zero phase lag) and low-pass filtered (100 Hz, 91 points, Hann window, zero phase lag), and were re-referenced to the average of the bilateral mastoid electrodes. The number of electrodes was reduced to 60 channels in order to reduce the size of data files. Independent components analysis was subsequently used to suppress artifacts due to eye movements, eye blinks, muscle movements, and heart beats (components were manually identified based on their topographical distributions and time course).

In three types of analyses (described below), we investigated whether language scores are related to the dynamics of the neural alpha oscillator: (1) alpha power, (2) alpha frequency flexibility, and (3) temporal correlations in alpha-amplitude time series.

### 2.4 Segmentation of data into epochs

For the alpha-power analysis and for the analysis of alpha-frequency flexibility, EEG data were segmented into six-hundred 7-second epochs. To this end, 7-second epochs were randomly selected from each 1-minute trial (three eyes-open trials and three eyes-closed trials) and were stored for subsequent analyses if the amplitude range in any of the electrodes did not exceed 300 μV (artifact rejection). This procedure was repeated until 100 artifact-free 7-second epochs were obtained for each 1-minute trial. This led to three hundred epochs for the eyes-open condition and three hundred epochs for the eyes-closed condition. The young age of the participants introduced infrequent artifacts in the data (e.g., movement), which, however, prohibited using the full 1-min data. The selection of 7-s data epochs provided sufficient artifact-free epochs without compromising the analysis approaches. Critically, calculation of power and frequency flexibility was carried out for each epoch independently (see below), and the resulting estimates were subsequently averaged across epochs. The use of multiple (even partially overlapping) data segments thus maximizes the amount of data used for the analysis, strengthens the average estimate, while reducing noise.

For the analysis of temporal correlations in alpha-amplitude time series, the EEG data were segmented into sixty 20-second epochs (separately for eyes-open and for eyes-closed conditions; i.e., 120 epochs overall). Analysis of temporal correlations requires a longer time course in order to capture neural dynamics accurately [25,48]. The choice of 20-second epochs was a compromise between the benefits of long epochs for this analysis and sufficient artifact-free data segments. Past studies demonstrated that temporal correlations in the EEG signal can be robustly detected with epochs as short as 20 seconds in duration [43,49,55]. A 20-second epoch was considered artifact-free if the standard deviation of any 200-ms segment of the epoch was smaller than 80 μV. For some participants (*n* = 8), there was an insufficient number of 20-second epochs that were artifact-free. Analysis of alpha-amplitude temporal correlations was thus performed for the remaining 33 participants (4 years, *n* = 8; 5 years, *n* = 14; 6 years, *n* = 11; Mean age: 5.6).

### 2.5 Alpha power analysis

For each 7-second epoch, a fast Fourier transform (FFT) was calculated (tapered by a Hann window; zero-padding) and the squared magnitude of the resulting complex values was used to obtain a power spectrum for each 7-second epoch. Power spectra of single 7-second epochs were averaged separately for the eyes-open and the eyes-closed conditions. The current analysis focused on the alpha frequency range (7–10 Hz). Note that the prominent frequency of the alpha oscillation is lower in children (~8.5 Hz) compared to adults (~10 Hz) [56–58]. Topographical distribution for the eyes-open and eyes-closed conditions was determined by calculating the mean power in the 7–10 Hz frequency band. In order to assess the difference between the eyes-open and eyes-closed conditions, the average power in the alpha frequency range was calculated across 10 posterior-occipital channels (Oz, O1, O2, PO3, PO4, P7, P8, P3, Pz, P4). The selection of channels was motivated by previous work showing that alpha power is strongest at parietal and occipital sites ([59–61]; also see Fig. 1C). The difference in alpha power between the eyes-open and eyes-closed conditions was assessed using a Wilcoxon signed-rank test.

**Figure 1.**
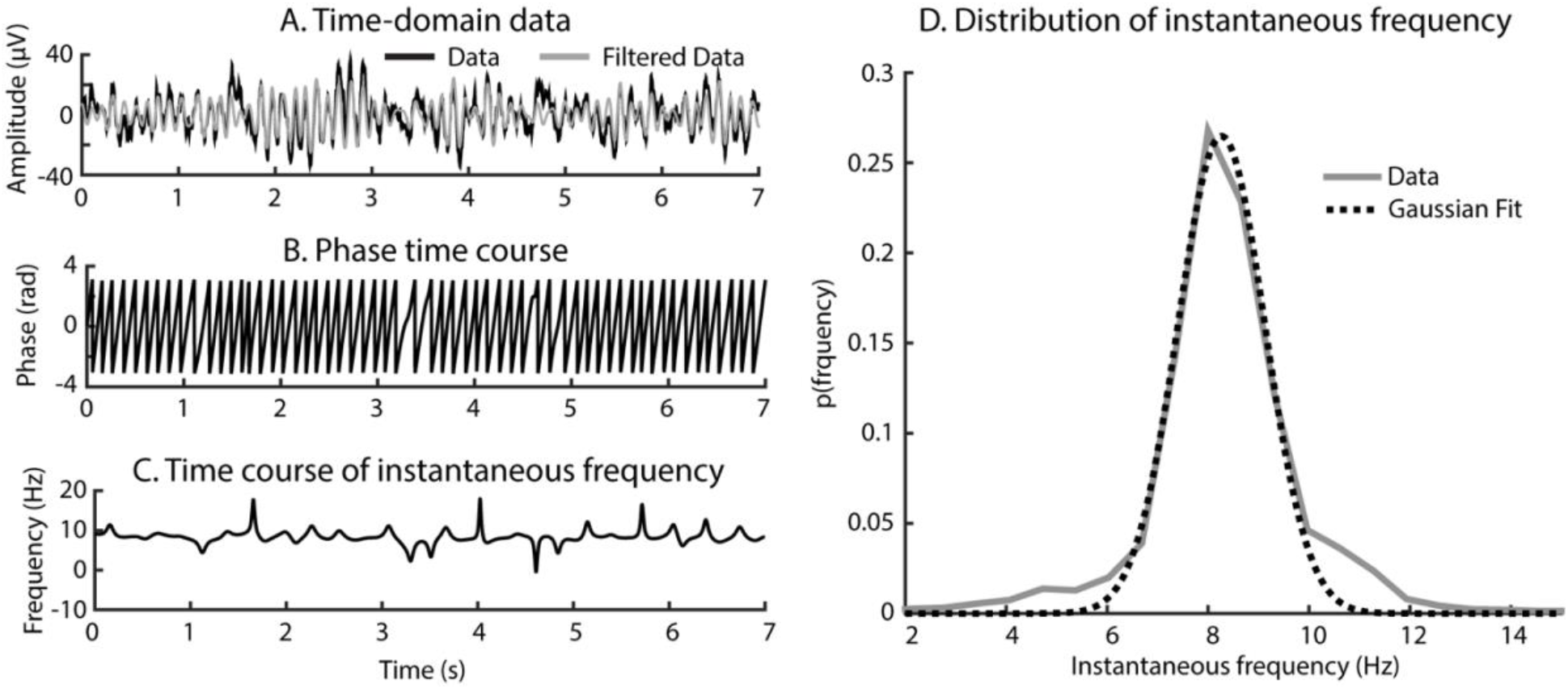
Estimation of the flexibility of the frequency of the neural alpha oscillator. **(*A*)** The original signal from one example epoch and its filtered version (5–12 Hz band-pass filter) are shown. **(*B*)** The phase angle (in rad) was calculated from the filtered signal (using Hilbert transformation). The changes in phase angle (y-axis) over time (x-axis) are plotted. Note that phase angles wrap at π and –π because angles are circular data. **(*C*)** The phase-angle time series was then transformed to a time series of instantaneous frequency. **(*D*)** Histogram of instantaneous frequency (i.e., the probability of a frequency [y-axis] as a function of frequency). A Gaussian function was fit to the histogram and the estimated width of the Gaussian function was used as a measure of the dynamic range of alpha frequency, and hence as a measure of frequency flexibility (a wider width reflecting a more flexible frequency).

A Spearman correlation was conducted in order to investigate the relation between alpha power difference (i.e., the difference between eyes-closed and eyes-open) and language ability. We also explored whether the relation between alpha power and language ability was independent of a participant’s chronological age and PIQ using a regression analysis with all three variables as predictors of alpha power (the difference in alpha power between eyes-closed and eyes-open was rank-transformed in order to account for the non-Gaussian distribution of power values; note that all results reported in this paper were also significant using parametric data analysis). The alpha power difference was highly correlated with the alpha power when children were resting with their eyes closed (*r* = 0.935) and open (*r* = 0.704).

### 2.6 Flexibility of the frequency of the alpha oscillator

Flexibility of the frequency of the alpha oscillator was analyzed using the 300 epochs for the eyes-closed conditions (7-second duration). We focused here only on the eyes-closed condition because estimation of alpha frequency requires the presence of alpha power (and alpha power is known to be greater when individuals have their eyes closed) and controlling for overall response magnitude, which is important for power, does not apply to frequency. Analysis of the flexibility of alpha frequency entailed the following steps (see Figure 1). Each 7-second epoch was filtered with a band-pass of 5–12 Hz (Kaiser window; centered on 8.5 Hz, see Figure 1A). The filter bandwidth was chosen to be wider here compared to the 7–10 Hz used for the alpha power analysis to avoid restricting the estimation of frequencies to a narrow range, and none of the results reported below were affected by adopting a different filter bandwidth (e.g., 3–14 Hz or 2–15 Hz). For each filtered epoch, the Hilbert transform was calculated (resulting in complex values) followed by calculation of the phase angle (Figure 1B). This resulted in a phase-angle time course for each epoch, which, in turn, was transformed to a time course of instantaneous frequency (Figure 1C). Individual steps of the analysis are depicted in Figure 1.

Quantification of the fluctuations in alpha frequency was calculated by binning the instantaneous frequency values into non-overlapping bins and calculating the probability of occurrence; this resulted in a histogram of instantaneous frequency values. A Gaussian function was fitted to the histogram of binned instantaneous frequency and the estimated width of the Gaussian function was taken as a measure of the dynamic range of alpha frequency, and hence as a measure of frequency flexibility (with a wider width reflecting a more flexible frequency; Figure 1D). The above steps were repeated for each channel, epoch, and participant. The width of the Gaussian function was averaged across the 10 posterior-occipital channels and across epochs. Note that for a few time points, the instantaneous frequency could not be estimated accurately (leading to very large or very small frequency values; Figure 1C) because the phase-angle from the Hilbert transform included rare jumps. However, our binning and Gaussian-fit approach is resistant to these inaccuracies because a few very small and very large instantaneous frequency values result in low probabilities at the tails of the histogram and thus do not affect the Gaussian fit.

In order to investigate the relation between the flexibility of alpha frequency and language ability, Spearman’s correlation was calculated between language scores and the estimated width of the Gaussian function in the eyes-closed condition. We also explored whether the relation between alpha frequency flexibility and language ability was independent of a participant’s chronological age and PIQ. To this end, we performed a regression analysis with age, PIQ, and language scores as predictors of rank-transformed alpha-frequency flexibility.

### 2.7 Long range temporal correlations in alpha-amplitude time series

Long-range temporal correlations in the alpha frequency band were analyzed using detrended fluctuation analysis [48,62–64]. Detrended fluctuation analysis was developed to measure irregular temporal correlation patterns [65], but has also been described in detail for its application to EEG oscillations [25,48]. Detrended fluctuation analysis is sensitive to the 1/*f* structure in non-stationary signals (that is, to the signal’s scaling behavior) and quantifies the long-term autocorrelation (for a tutorial on detrended fluctuation analysis see [25]).

For the current investigation, the detrended fluctuation analysis involved the following steps (depicted in Figure 2). Each 20-second epoch was band-pass filtered at 7–10 Hz in order to extract the alpha frequency band. The alpha-amplitude envelope was estimated by calculating the Hilbert transform of the filtered signal, followed by computing the magnitude of the resulting complex values. The mean across the time points of the 20-second amplitude envelope was subtracted from each of the amplitude envelope’s time points (Figure 2A), and the cumulative sum was calculated for the mean-centered amplitude envelope (Figure 2B). Sliding windows (*N* = 12) of different size were subsequently shifted across the amplitude-envelope time series (50% overlap). Window sizes were logarithmically spaced and ranged from 1 to 19.5 seconds. For each shift, the signal was detrended using linear regression (i.e., slope and intercept were removed; Figure 2C) and the root-mean-square amplitude was calculated for the detrended signal. The root-mean-square amplitude was averaged across all shifts of a particular window size. This resulted in a root-mean-square value for each window size, which when displayed on logarithmic axes shows a linear trend (Figure 2D). A linear function was fit to the log-transformed root-mean square values as a function of the log-transformed window size. The estimated slope (i.e., beta coefficient) of the fitted linear function reflects the degree of long-range correlation in the alpha-amplitude time course. The beta coefficient was calculated for each 20-second epoch, electrode, condition (eyes-open, eyes-closed), and participant. Beta coefficients were averaged across the sixty 20-second epochs (independent for eyes-open and eyes-closed) and across the 10 posterior-occipital electrodes.

**Figure 2.**
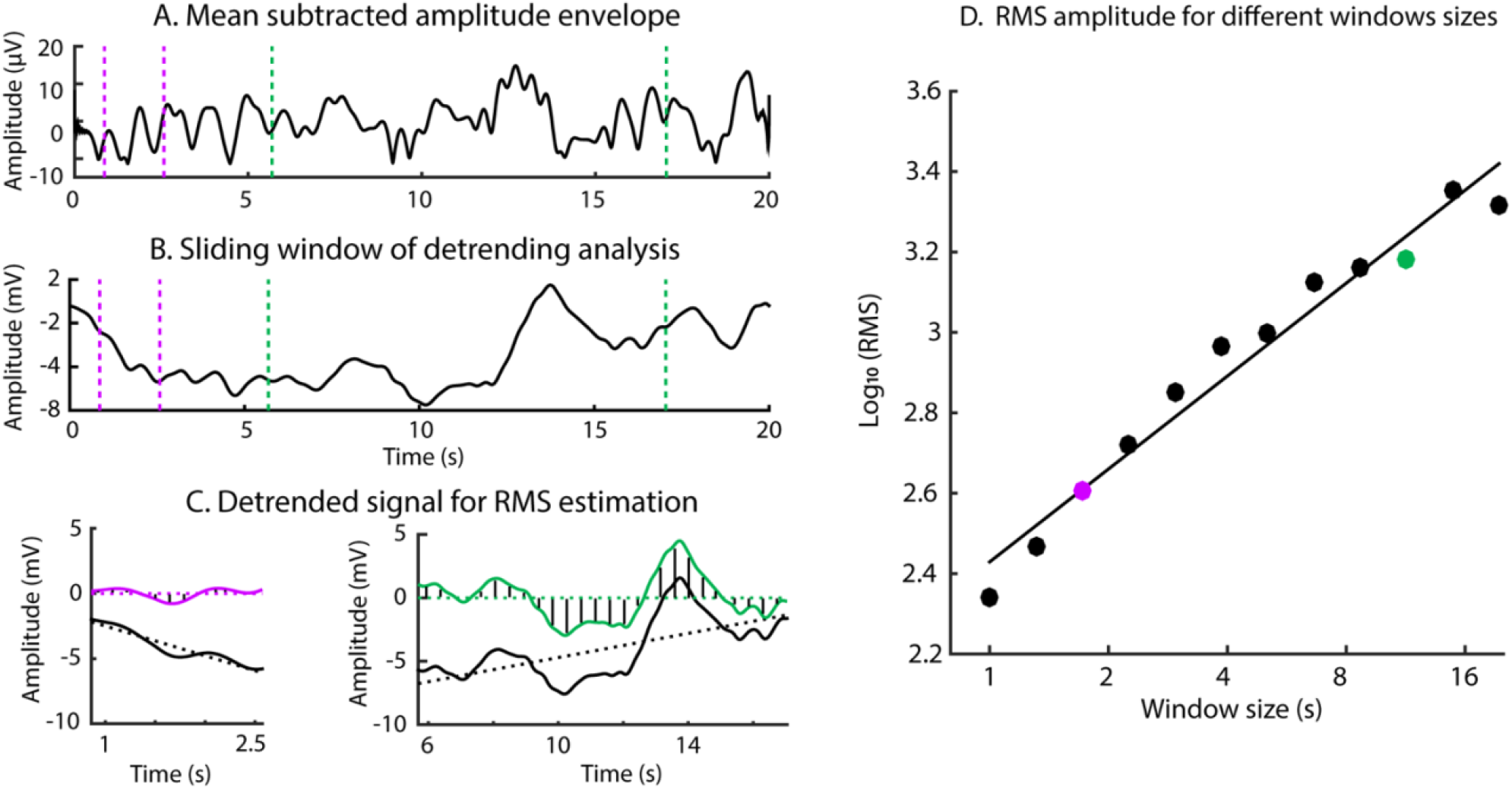
Calculation of the detrended fluctuation analysis for the alpha-amplitude time series. **(*A*)** The alpha-amplitude envelope (with mean subtracted) for an example epoch taken from the eyes-closed condition of one participant. **(*B*)** The cumulative sum of the signal in panel A. The dashed, vertical lines mark two example time windows (1.7 seconds and 11.4 seconds). **(*C*)** Signals (black solid lines) from the two time windows depicted in B (left 1.7 seconds; right 11.4 seconds). Detrended signals are displayed in color. The dashed lines reflect the linear regression line that was subtracted from the signal to obtain the detrended signal. The root-mean-square (RMS) amplitude of the detrended signal was calculated (marked by the lines that cover the area under the detrended signal), which is a measure of amplitude variation. **(*D*)** The log value of the RMS (y-axis) is plotted against each of the window lengths analyzed (x-axis). Colored dots in this panel mark the RMS calculated for signals depicted in panel C.

The detrended fluctuation analysis is sensitive to the relative degree of energy in different neural frequency bands and thus to the 1/*f* structure in the alpha-amplitude time series (see also [25]). For example, a large beta coefficient (steep slope) indicates that the fluctuations in signal (as estimated using the root-mean-square) are relatively large for wide window sizes and relatively small for narrow window sizes. In this case, energy in low-frequency bands (i.e., wide windows) is high and the signal therefore contains long-term autocorrelations. In contrast, a small beta coefficient (shallow slope) indicates that the fluctuations in signal are relatively small for wide window sizes and relatively large for narrow window sizes. In this case, energy in low-frequency bands (i.e., wide windows) is low and the signal therefore contains less long-term autocorrelations.

In order to quantify whether significant long-range correlations were present in the alpha amplitude time series, we calculated the detrended fluctuation analysis for surrogate data and subtracted the estimated beta coefficients of the surrogate data from the beta coefficients of the original time series. Surrogate data were obtained by calculating a fast Fourier transform for each 20-s epoch, assigning a random phase value for each frequency bin (while keeping the amplitude information), and inverting the Fourier transform. This procedure provides a time series in which the temporal organization is more random compared to the original time series. Calculation of surrogate data and computation of detrended fluctuation analysis was repeated 20 times. The 20 estimated beta coefficients from the detrended fluctuation analysis on surrogate data were averaged. The difference in beta coefficient between the eyes-open and eyes-closed conditions, and the difference between the beta coefficients and the surrogate-beta was assessed using a Wilcoxon signed-rank test.

Finally, we explored the relations between the different properties of the alpha oscillation (i.e. power, frequency flexibility, and temporal correlation) using Spearman correlation.

## 3. Results

### 3.1 Alpha spectral power

Figure 3 depicts the power spectrum for the eyes-open and eye-closed conditions. Alpha power was significantly larger in the eyes-closed compared to eyes-open condition (Median = 6.67, 1.32 μV^2^ respectively, *p* < 0.001, *N* = 41; Figures 3A-C). The EEG topographies (Figure 3D) show that the changes in alpha power were strongest at the occipital and parietal regions of the scalp.

**Figure 3.**
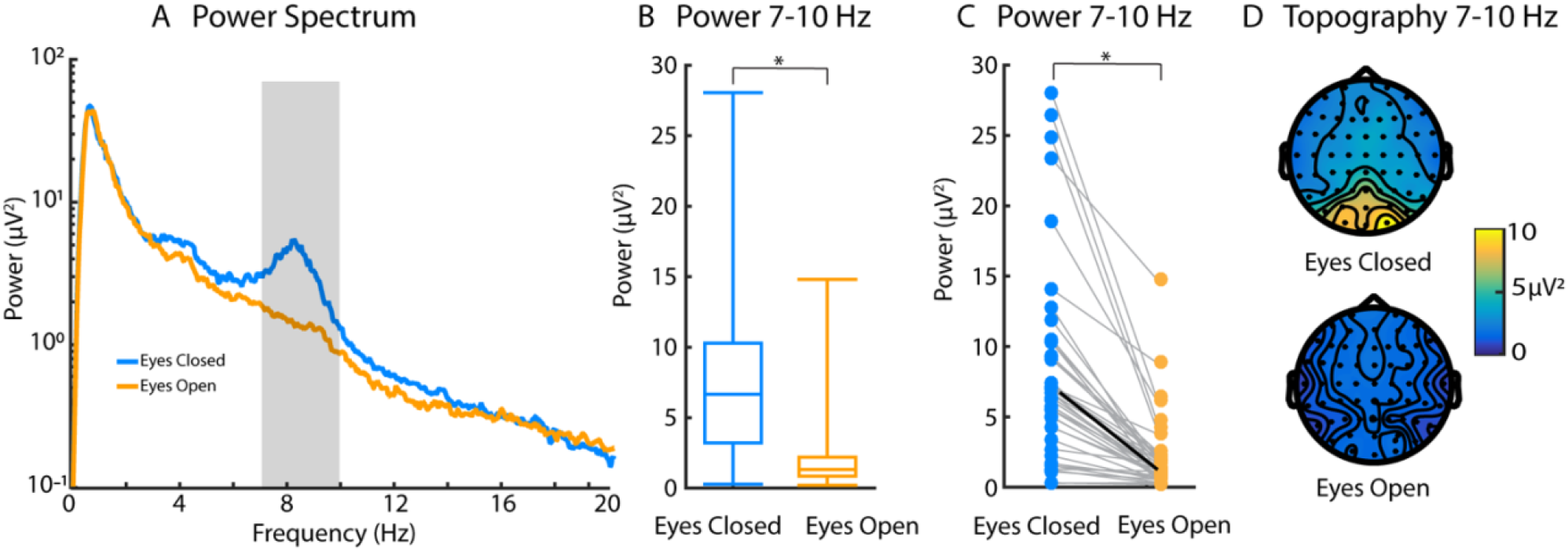
Alpha power analysis. **(A)** Power spectra for the eyes-open and eyes-closed condition. The gray area marks the alpha-frequency range 7–10 Hz. **(B)** Box plots show the alpha power for each condition for the 7–10 Hz range (eyes-open, eyes-closed). **(C)** Same data as in panel B, here showing the alpha power for each child. **(D)** Scalp distributions of alpha power (7–10 Hz). \**p* < 0.001

A significant negative correlation was found between the difference in alpha power (∆alpha-power; eyes-closed minus eye-open conditions) and language abilities (*r* = –0.363, *p* = 0.022; Figure 4, also summarized in Table 1). The regression including chronological age, PIQ in addition to language as predictors for the rank-transformed ∆alpha-power was not significant (*p* = 0.143), although language ability remained a significant predictor when considered in isolation (*t*(37) = –2.164, *p* = 0.037). A summary of the regression analysis is provided in Table 2.

**Figure 4.**
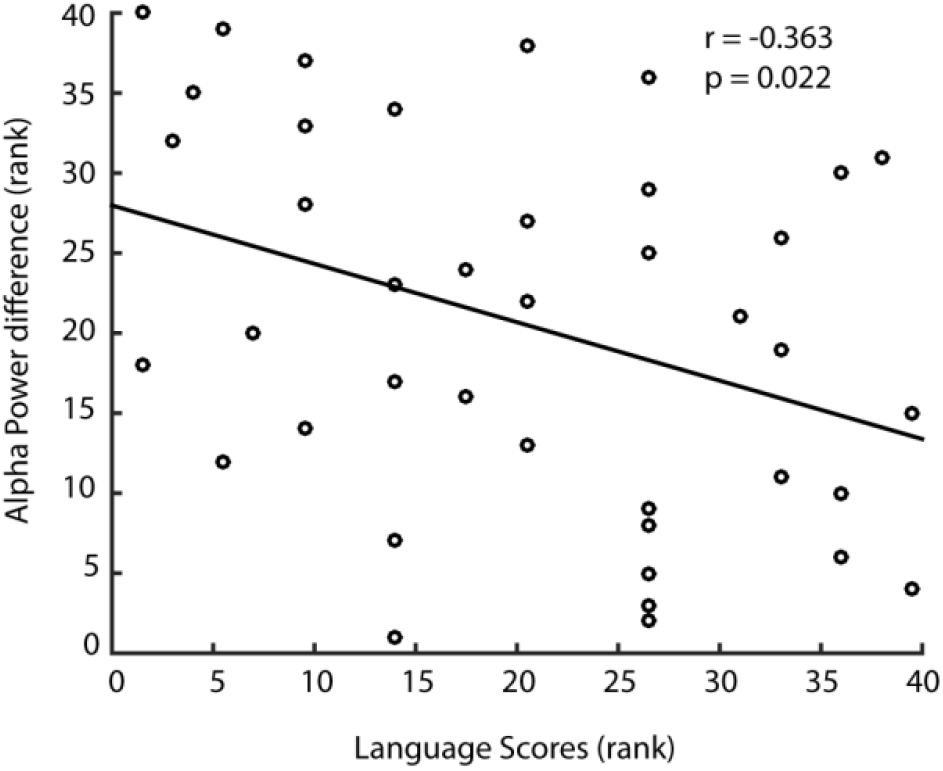
Spearman’ correlation between language score and alpha power difference (eyes-closed minus eyes-open).

**Table 1.** Correlation Matrix Between Properties of Alpha Oscillation and Language Abilities

| | Language<br>ability | Δalpha-power | Alpha-frequency<br>dynamics | Δalpha-temporal<br>correlation |
| --- | --- | --- | --- | --- |
| Language ability |  |  |  |  |
| Δalpha-power | $r = -0.363$<br>$p = 0.022^*$ | | | |
| Alpha-frequency<br>dynamics | $r = 0.426$<br>$p = 0.006^*$ | $r = -0.643$<br>$p < 0.001^*$ | | |
| Δalpha-temporal<br>correlation | $r = 0.480$<br>$p = 0.005^*$ | $r = -0.205$<br>$p = 0.253$ | $r = 0.100$<br>$p = 0.584$ | |

**Table 2.** Summary of Regression Analysis on Properties of Alpha Oscillation

| Alpha properties | *t* | *p* | $\beta$ | *F* | *df* | *p* | *R*<sup>2</sup> |
| --- | --- | --- | --- | --- | --- | --- | --- |
| Δalpha-power | | | | 1.923 | 3,37 | 0.143 | 0.138 |
| Chronological age | 0.276 | 0.784 | 0.582 |  |  |  |  |
| Performance IQ | -0.094 | 0.925 | -0.016 |  |  |  |  |
| Language Score | -2.164 | 0.037* | -0.464 |  |  |  |  |
| alpha frequency dynamics |  |  |  | 3.772 | 3,37 | 0.019* | 0.239 |
| Chronological age | -0.175 | 0.862 | -0.347 |  |  |  |  |
| Performance IQ | -1.726 | 0.093 | -0.269 |  |  |  |  |
| Language Score | 3.237 | 0.003* | 0.652 |  |  |  |  |
| Δalpha-temporal correlation | | | | 4.596 | 3,29 | 0.010* | 0.322 |
| Chronological age | -0.519 | 0.608 | -0.975 |  |  |  |  |
| Performance IQ | -2.311 | 0.028* | -0.357 |  |  |  |  |
| Language Score | 3.624 | 0.001* | 0.672 |  |  |  |  |

\* $p < 0.05$
Note: We also conducted additional regression analyses (not reported here) to confirm that all properties of alpha oscillation were significant *predictors* (in contrast to dependent measures) of language ability after controlling for the variance contributed by the performance IQ and chronological age.

### 3.2 Flexibility of alpha frequency

The flexibility of the alpha oscillator’s frequency was estimated as the width of a Gaussian function that was fit to binned instantaneous neural frequency values (see Methods). A significant correlation was found between the flexibility of alpha-frequency (i.e., the estimated Gaussian width using the eyes-closed condition) and language abilities (*r* = 0.426, *p* = 0.006; Figure 5, also summarized in Table 1). The overall regression analysis considering age, PIQ, and language score as predictors for the rank-transformed alpha-frequency flexibility was significant (*R*^2^ = 0.239, *p* = 0.019). However, only language score (*ß* = 0.652, *t*(37) = 3.237, *p* = 0.003) was a significant predictor of the flexibility of alpha-frequency (results summarized in Table 2).

**Figure 5.**
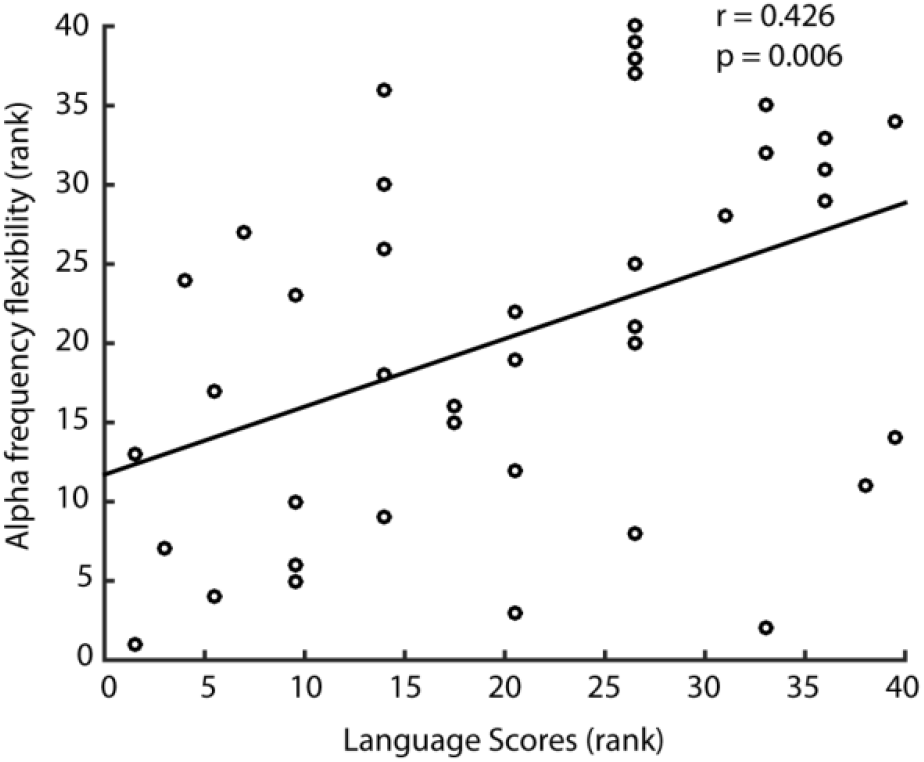
Spearman’ correlation between flexibility of alpha-frequency (estimated width of Gaussian fit) and language ability.

### 3.3 Long-range temporal correlation of alpha oscillation

The long-range temporal correlations in the alpha amplitude time series was analyzed using the beta coefficient from the detrended fluctuation analysis. For the original beta coefficients (non-corrected by surrogate data), the coefficient was larger in the eyes-closed compared to the eyes-open condition (*p* = 0.001; *N* = 33). The original beta coefficient was also larger compared to the beta coefficient from surrogate data for the eyes-closed (*p* < 0.001) and the eyes-open (*p* < 0.001) conditions, indicating that long-range temporal correlation were present in the alpha amplitude time series. No difference between eye-closed and eyes-open was found for beta coefficients corrected using surrogate data (*p* = 0.416, *N* = 33; Figure 6B, 6C). The EEG topography (Figure 6A) shows the largest beta coefficients at occipital electrodes.

A significant correlation was found between the Δalpha-temporal correlation (i.e., difference in surrogate-data-corrected beta coefficients between the eyes-closed and eyes-open condition) and language score (*r* = 0.480, *p* = 0.005; Figure 6D, also summarized in Table 1). The overall regression analysis considering age, PIQ, and language score was significant (*R*^2^ = 0.322, *p* = 0.010). ∆alpha-temporal correlation (rank-transformed) was predicted by both the children’s language abilities (*ß* = 0.672, *t*(29) = 3.624, *p* = 0.001) and PIQ (*ß* = −0.357, *t*(29) = −2.311, *p* = 0.028). ∆alpha-temporal correlation was not predicted by children’s chronological age (also summarized in Table 2). Note that language scores also predicted ∆alpha-temporal correlation (difference in beta coefficients) in a regression analysis using the non-corrected beta coefficients (i.e., without correction by surrogate data) (*t*(29) = 2.265; *p* = 0.031).

**Figure 6.**
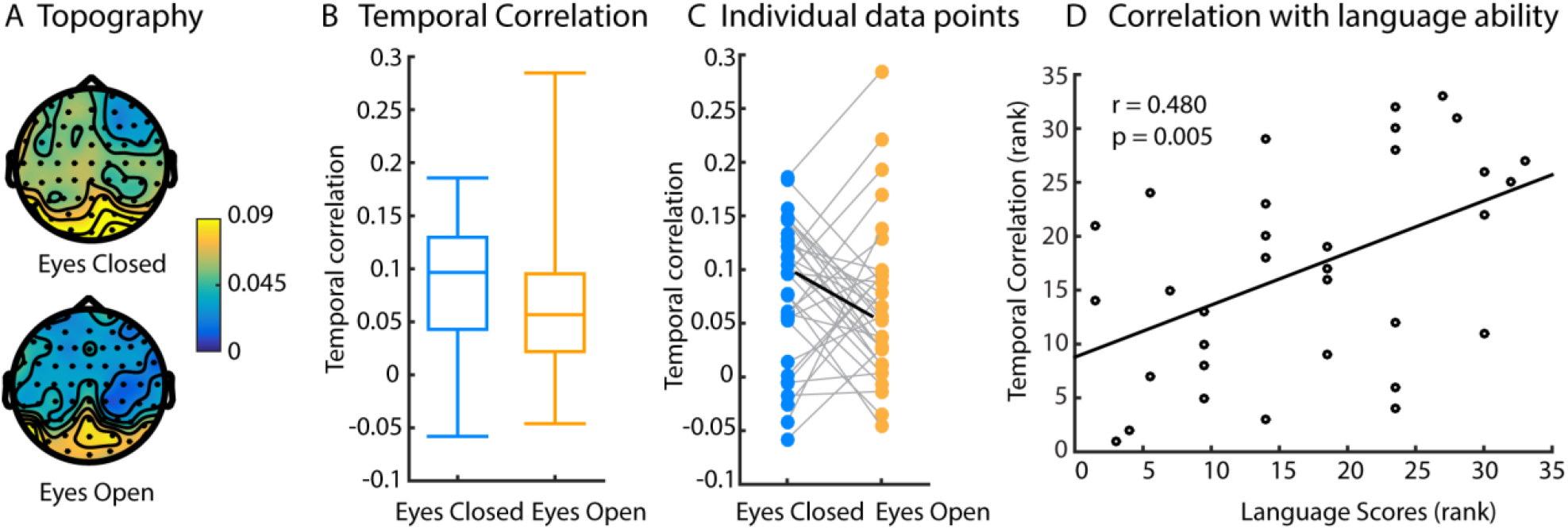
Analysis of long-range temporal correlations in alpha amplitude time series. **(A)** Scalp distribution of beta coefficients from detrended fluctuation analysis that indicate the degree of long-range correlations; larger values index greater long-range correlations. **(B)** Box plots show the beta coefficients (corrected using surrogate data) for each condition. **(C)** Same data as in panel B, here showing the beta coefficient for each child. **(D)** The differences in beta coefficients between conditions (eyes-closed minus eyes-open) was positively correlated with language scores.

### 3.4 Relation between properties of alpha activity

The only significant relation found between the different indexes of alpha activity was between alpha power and frequency flexibility. The difference in alpha power between the eyes-closed and eyes-open conditions was negatively correlated with frequency flexibility (*r* = −0.643, *p* < 0.001, also see summary in Table 1). Hence, when alpha frequency is widely fluctuating across frequencies, alpha power tends to be smaller. This may not be surprising as power (measured in the static power spectrum) will be more spread out across frequencies in the case of high frequency flexibility. Critically, although the two measures are correlated, they only share about 41% of variance. Both measures may thus reflect useful indices to describe spontaneous activity. Temporal correlation of alpha oscillations was not correlated with alpha power or frequency flexibility.

## 4. Discussion

In the current study, we investigated the relation between oral language ability and resting-state neural activity in typically developing children aged 4 to 6 years. Our main goal was to explore properties of the neural alpha oscillator that have been largely overlooked in the language literature. We found that the power, frequency flexibility, and temporal correlation of the alpha oscillation correlated with children’s performance on a standardized language measure. These findings suggest that a comprehensive examination of neural oscillator properties with multiple analytic techniques may be required to reveal the role of neural oscillations in cognition.

### 4.1 Oscillation power and oral language

We found a negative correlation between oral language ability and resting-state alpha power. That is, the alpha power difference between eyes-closed and eyes-open during rest was smaller for children who had the most advanced language skills relative to their same-age peers. This relation between alpha power and spoken language ability is consistent with some of the literature on reading disorder populations, where elevated resting alpha power has been noted in school-age and preschool children with reading disorders [18,27,66]. Our regression analysis further demonstrates that the relation between alpha power and oral language ability may be influenced by chronological age and performance IQ, as a significant model was not found after controlling for the variance contributed by these variables.

A prominent theory suggests that alpha oscillations reflect a mechanism that gates inhibition in neural circuits [13,67]. Under this view, a low level of alpha activity may indicate a higher level of cortical excitability, which impacts cognitive functions [68,69]. While our study did not investigate the role of resting alpha oscillation power in sensory processes, previous work has found evidence to support a role of alpha oscillations in sensory processing. For example, the ability to detect near-threshold stimuli is reduced when alpha power immediately before the stimulus presentation is high (e.g. visual stimuli: [70–74]; e.g. somatosensory stimuli: [75,76]; e.g. auditory stimuli [77]), but more complex relations between alpha power and perception have been reported as well [78,79].

The extent to which alpha power immediately before stimulus presentation is related to observations of alpha power when an individual is at rest (i.e., EEG recorded in the absence of sensory stimulation over several minutes) will warrant further investigation. Previous work found that alpha power at rest was different between good and poor performers on sensory detection tasks. Hanslmayr et al. (2007) reported that *perceivers* (those who detected visual stimuli at a higher than chance level) have significantly lower alpha power at rest compared to *non-perceivers* (those who detected visual stimuli at chance level). The authors suggested that this difference in spontaneous alpha power underlies the pre-stimulus alpha power difference in detected versus missed trials. In future studies, it may be useful to explore the relation between alpha power and performance on sensory detection tasks as the mediator of the relation between alpha power and spoken language ability.

Beyond alpha, spontaneous frontal gamma activity (31–50 Hz) has also been found to not only significantly correlate with concurrent oral language skills [23] but also to later language skills [24]. In the current study, our analysis focused on the occipital alpha oscillation as its role in oral language ability is unclear. In addition, occipital alpha oscillation dominates the EEG activity at rest, and thus provides a better signal-to-noise ratio for our EEG analysis approaches. The estimation of long-range temporal correlation may be challenging for neural frequency bands such as gamma, which show no clear peak and have low signal-to-noise ratios [25]. The presence of an identifiable peak in the power spectrum (see Figure 3A) suggests the existence of an underlying neural oscillator (here in the alpha range), whereas activity without a clear peak in the power spectrum may indicate non-rhythmic rather than oscillatory activity [39,40]. Furthermore, previous work consistently demonstrated that alpha oscillations play a significant role in cognitive functions [65,78–80], which made it a good candidate in our current exploratory study.

### 4.2 Dynamics of alpha oscillation: Frequency flexibility and temporal correlations

In typically developing children, we found a positive correlation between oral language ability and the frequency flexibility of the alpha oscillation. That is, children with an alpha oscillator that exhibits higher flexibility had more sophisticated spoken language abilities relative to their peers. Furthermore, we observed that greater long-range temporal correlations in the alpha-amplitude time series were associated with higher language scores. Our analyses of alpha frequency flexibility and long-range temporal correlations consider the moment-to-moment fluctuations in EEG oscillations that are often dismissed as noise, but may be important for cognition [83,84].

Our results corroborate the expanding literature that demonstrates the existence of critical-state dynamics in human neural systems [47,48,85–87]. At criticality, neural networks perform at maximal variability while maintaining stability [47,88]. Computational models and empirical studies suggest several advantages for neural networks to function at a critical state, including maximal information transmission and storage [35,88–90], as well as an optimized dynamic range that fosters stimulus processing [88,91,92]. Furthermore, cognitive functions may be optimized when neural activity operates near criticality, benefiting, for example, the range of stimuli distinguishable by a cognitive network, the fidelity of information being transmitted, and the capacity of a system to engage in different tasks [88]. In our current analysis, we investigated two aspects of criticality: frequency flexibility and long-range temporal correlation, both of which were positively correlated with language ability.

Variability in network activation patterns at criticality has been shown to enhance the detection of weak stimuli [93,94] and to prevent degradation of sensory information during neuronal communication [95]. In terms of the strength of temporal correlation, previous work suggests that greater temporal (i.e., auto-) correlation in time series may reflect the ability of the brain to sustain a stable oscillation over time for cognitive processing, such as in the case of working memory tasks [62,96]. A reduction in temporal correlation, which has been observed in patients with Alzheimer’s disease and healthy individuals who are sleep deprived, may underlie the reduced cognitive flexibility [97,98] and poorer ability to integrate information [99] observed in these individuals. One study of young adolescents (sixth to eighth grade) found that those with developmental reading disability had reduced temporal organization in resting MEG recordings compared to same-age peers, thus suggesting a role of self-organized criticality in language development [100]. Findings from our study add further support for the role of long-range temporal correlation in cognitive functions.

Considered together, our findings suggest that children who have more advanced language abilities relative to their same-age peers may have neural systems that function closer to criticality, such that the neural alpha oscillator flexibly alters its natural frequency and temporal correlations are maximally maintained in its time course.

### 4.3 Limitations and future directions

We demonstrated that the power, flexibility of the alpha frequency, and long-range temporal correlation of resting alpha oscillations correlated with oral language ability. Given the exploratory nature of our approach, several additional questions remain worthy of further examination. One future direction is to clarify the cognitive mechanisms that relate resting-state alpha oscillation to language ability. Our work, and work from other groups, suggests that immature processing of auditory stimuli, including those presented in rapid succession, is related to language disorders [101–104], particularly impairment in receptive language [104–106]. It will be important to investigate whether auditory perception mediates the relation between alpha oscillatory activity and language ability. In the current study, we observed a moderate correlation between properties of the alpha oscillator and language ability in a population of children with typical development. The extent to which properties of the neural alpha oscillator predict disordered language development, or impairment in different domains of oral language (such as expressive versus receptive), is unknown, but our data may provide a fruitful starting point for the exploration of neural markers for language disorder.

Developmentally, a power-law structure in the power spectrum of resting EEG recordings can be found in newborns [107], suggesting that brain development toward criticality occurs very early in life, and continues to mature into adulthood and beyond [62]. What we do not yet fully understand is the relation between the different properties of neural oscillators (in particular across the lifespan). Previous studies have shown that long-range temporal correlations in the alpha time series are not correlated with power [55], and display a maturational trajectory that is independent of alpha power [62]. We also found that temporal correlation of alpha activities did not correlate with alpha power in our data, which suggests a distinct developmental role for different properties of the alpha oscillator. There is a need for future research, particularly longitudinal research, to explore beyond the traditional properties of neural oscillators (i.e., power spectrum) in order to reveal the intricate roles of neural oscillators in oral language ability and other cognitive processes.

## Acknowledgements

This work was supported by Natural Science and Engineering Research Council of Canada and Ontario Research Fund. BH was supported by a BrainsCAN postdoctoral fellowship (Canada First Research Excellence Fund; CFREF). There are no conflicts of interest associated with this study.

